# An efficient deep learning method for amino acid substitution model selection

**DOI:** 10.1101/2024.06.27.600948

**Authors:** Nguyen Huy Tinh, Le Sy Vinh

## Abstract

Amino acid substitution models play an important role in studying the evolutionary relationships among species from protein sequences. The amino acid substitution model consists of a large number of parameters; therefore, it is estimated from hundreds or thousands of alignments. Both general models and clade–specific models have been estimated and widely used in phylogenetic analyses. The maximum likelihood method is normally used to select the best fit model for a specific protein alignment under the study. A number of studies have discussed theoretical concerns as well as computational burden of the maximum likelihood methods in model selection. Recently, machine learning methods have been proposed for selecting nucleotide models. In this paper, we propose methods to create summary statistics from protein alignments to efficiently train a network of so-called ModelDetector based on the convolutional neural network ResNet-18 for detecting amino acid models. Experiments on simulation data showed that the accuracy of ModelDetector was comparable with that of the maximum likelihood method ModelFinder. The ModelDetector network was trained from 64,800 alignments on a computer with 8 cores (without GPU) in about 12 hours. It is orders of magnitudes faster than the maximum likelihood method in inferring amino acid substitution models and able to analyze genome alignments with million sites in minutes.

## Introduction

Identifying proper amino acid (AA) substitution models for analyzing protein alignments under the study is one of the fundamental steps in studying the evolutionary relationships among species from their protein data. The amino acid substitution models are normally assumed to be time-reversible, time-homogeneous, time-continuous and stationary (Tavaré 1986; Felsenstein 2003). An amino acid substitution model can be represented by a 20 × 20 substitution rate matrix *Q* = {*q*_*xy*_} where coefficient *q*_*xy*_ represents the substitution rate from amino acid *x* to amino acid *y*. Up to date, a number of amino acid substitution models have been introduced including widely-used general models estimated from large diverse datasets, e.g., JTT (Jones, Taylor, and Thornton 1992), WAG (Whelan and Goldman 2001), LG (Le and Gascuel 2008), Q.pfam (Minh et al. 2021); and clade-specific models each estimated from proteins of the same clade such as Q.plant, Q.bird, Q.yeast, Q.mammal, Q.insect (Minh et al. 2021).

The maximum likelihood method has been widely used to select the best fit model for a specific alignment. It uses information criteria such as BIC (Schwarz 2007) or AIC (Akaike 1974) for determining the best fit model, e.g., ModelFinder (Kalyaanamoorthy et al. 2017) or ModelTest-NG (Darriba et al. 2020). However, using information criteria for model selection is still questionable in phylogenetic analyses (Crotty and Holland 2022; Grievink et al. 2010; Jhwueng et al. 2014; Seo and Thorne 2018; Susko and Roger 2020; Burgstaller-Muehlbacher et al. 2023; Nguyen Huy, Dang, and Sy Vinh 2023). The true trees and true models might not have the best BIC/AIC values. Besides theoretical concerns, finding the best fit model by the maximum likelihood method is computationally prohibitive for large alignments because we have to construct trees and calculate their likelihood values with various available models. The model selection is one of the most computationally expensive steps in the maximum likelihood algorithms to partition alignments such as mPartition (Le Kim and Le Sy 2020).

Machine learning approach has been applied to solve a number of problems in biology (Kan 2017; Kandoi, Acencio, and Lemke 2015; Leung et al. 2016). It is a promising approach for the model selection problem, e.g., ModelTeller (Abadi et al. 2020) and ModelRevelator (Burgstaller-Muehlbacher et al. 2023). ModelTeller and ModelRevelator work well on simulation nucleotide data with high accuracy, e.g., 91% for GTR model (Burgstaller-Muehlbacher et al. 2023). As sequence alignments cannot be directly used as inputs for training deep learning networks, ModelRevelator creates a total of 260,000 summary statistics for each nucleotide alignment to train a convolutional neural network ResNet-18 (He et al. 2016). Training a ResNet-18 network with a huge number of summary statistics requires powerful machines with GPU hardware.

The amino acid substitution models consist of many more parameters than the nucleotide substitution models. In this paper, we introduce Pairwise and Triplet extraction methods each creates 400 summary statistics from protein alignments. The Pairwise extraction method creates 400 summary statistics from pairwise sequences that effectively captures information from closely related sequences. The Triplet extraction method calculates 400 additional summary statistics from triplet sequences that might handle long-distance sequences.

We trained two deep learning networks: pModelDetector network with 400 summary statistics calculated from the Pairwise extraction method and ModelDetector network with 800 summary statistics combined from both Pairwise and Triplet extraction methods. The networks were trained on simulation data generated from real alignments to recognize nine models: five clade-specific models, i.e., Q.plant, Q.bird, Q.yeast, Q.mammal and Q.insect (Minh et al. 2021); and four widely used general models, i.e., Q.pfam (Minh et al. 2021), LG (Le and Gascuel 2008), WAG (Whelan and Goldman 2001) and JTT (Jones, Taylor, and Thornton 1992). The performance of the deep learning networks were compared with that of the maximum likelihood method ModelFinder (Kalyaanamoorthy et al. 2017) on simulation data.

## Materials and methods

### Data

#### Data simulation

We simulated protein alignments from real alignments using the AliSim program (Ly-Trong et al. 2022). First, we simulated alignments with five clade-specific models: Q.plant, Q.bird, Q.yeast, Q.mammal and Q.insect (Minh et al. 2021). For each clade, we randomly selected 1000 real protein alignments (each alignment has at least 50 variant sites) available at https://github.com/roblanf/BenchmarkAlignments. In this study, we trained our networks with time reversible models so outgroup sequences were removed from the real alignments because they are unrelated to ingroup sequences. Table 1 shows data statistics of the real clade-specific alignments. For each real alignment *D* of a clade C, we used AliSim to simulate alignments with model Q.C (e.g., Q.plant for real plant alignments). The AliSim program was conducted with +G (gamma rate distribution model) and +I (invariant rate model) options to simulate alignments with site rate heterogeneity models. The tree and parameters of the site rate models used to simulate data were estimated from the real alignment *D*.

**Table 1.** The data statistics of real alignments from five clades. #MSA: the number of alignments; #Sites: the average number of sites; Branch length: the average branch length of trees.

| Clades | #MSA | #Sequences | #Sites | Branch length |
| --- | --- | --- | --- | --- |
| Plant | 1000 | 35 | 330 | 0.064 |
| Bird | 1000 | 48 | 676 | 0.018 |
| Yeast | 1000 | 332 | 508 | 0.169 |
| Mammal | 1000 | 82 | 689 | 0.027 |
| Insect | 1000 | 134 | 254 | 0.157 |

Second, we simulated alignments with general models. To this end, we downloaded 1000 general real alignments (each alignment has at least 30 sequences and 50 variant sites) from the HSSP database (Sander and Schneider 1994). For each general model *Q* (i.e., Q.pfam, LG, WAG, and JTT), we also used AliSim with +G and +I options to simulate alignments with *Q*. Trees and parameters of the site rate models used to generate simulation alignments were estimated from the real alignments.

The Fig. 1 shows data statistics of the 1000 real HSSP alignments. The average tree branch length of majority alignments ranges from 0.08 to 0.14 (only 5 alignments with average tree branch lengths higher than 0.25). More than half (57%) of alignments have less than 50 sequences; and only ∼6% of alignments with more than 150 sequences. Almost all alignments have at least 100 sites (only ∼3% of alignments with less than 100 sites).

**Fig. 1.**
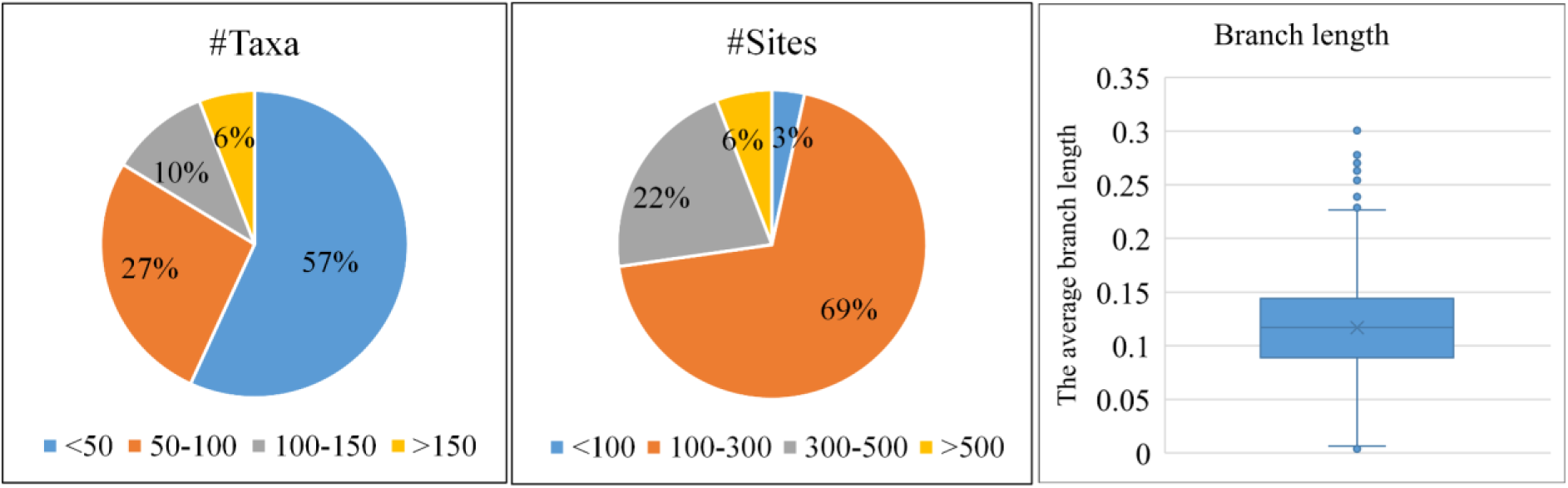
The distribution of average tree branch length, the number of taxa, and the number of sites of 1000 real HSSP alignments.

#### Training and testing data

We employed AliSim to simulate training and testing alignments from real alignments. Each real dataset (i.e., a clade-specific dataset or HSSP dataset) were split into a training part of 900 real alignments used to simulate training alignments and a testing part including the rest 100 real alignments employed to generate testing alignments. Technically, for each model we used the real alignments in the training part to simulate 7200 training alignments; and the real alignments in the testing part to generate 500 testing alignments. The simulation alignments were generated with different lengths (100, 500, 1000, and 2000 amino acids). In total, we generated 64,800 training alignments and 4500 testing alignments for 9 models. We used the training strategy that allocated 80% of the training alignments for training the networks and the remaining 20% of training alignments for the network validation.

### Methods

#### Maximum likelihood method

Given an alignment and a set of amino acid substitution models **Q**, the maximum likelihood model selection method will determine a tree *T* and a model *Q* ∈ ***Q*** to maximize the likelihood value *L*(*Q*, *T* | *D*). As models might have different number of parameters, information criteria such as BIC score (Schwarz 2007) can be used to select the best fit model.

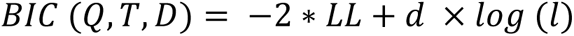

where *LL* is the log likelihood value of *L*(*Q*, *T*|*D*); *d* is the number of free parameters; and *l* is the alignment length. The lower BIC score indicates better substitution model.

Searching tree *T* and substitution model *Q* simultaneously is a computational challenge and onl y feasible for small alignments. A number of heuristic methods have been proposed to reduce the computational burden such as ModelFinder (Kalyaanamoorthy et al. 2017). ModelFinder selects the best fit model *Q* based on a quickly constructed tree *T*. For a protein alignment, ModelFinder approximately search the maximum likelihood tree *T* using the general model LG. Having the tree *T,* it selects the best fit model *Q*^∗^ ∈ ***Q*** that maximizes the BIC score

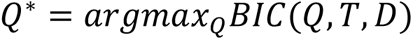

The models for site rate heterogeneity such as gamma distribution model with four rate categories (G4) and invariant site model (I) can be incorporated in the model selection process.

#### Deep learning methods

##### Pairwise extraction method

Creating summary statistics from protein alignments is one of the most crucial steps in training deep learning models for amino acid model selection. The *Q* matrix of an amino acid substitution model has 400 relative rates of change. For each protein alignment, we randomly select 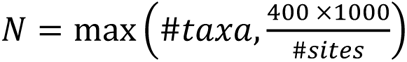 sequence pairs where #*taxa* and #*sites* are the number of sequences and alignment length, respectively. Amino acid substitutions from a pair of sequences does not contain enough information for all values of *Q*, therefore, amino acid substitutions in all pairs of sequences are counted into a 20 × 20 frequency matrix *F*_2_ = {*f*_2_(*xy*)} where *f*_2_(*xy*) is the number of times two amino acids *x* and *y* occur at the same position in sequence pairs. For each alignment, 400 relative rates of change of the *F*_2_ matrix summarizing amino acid substitutions from the alignment are used for training deep learning models.

##### Triplet extraction method

The genetic distance between two sequences of a pair might be large that might make the summary statistics of *F*_2_ less reliable. We introduce the Triplet extraction method to count amino acid substitutions from three sequences to generate 400 other summary statistics. Let *S*_1_, *S*_2_, and *S*_3_ be three different sequences; and *C* be their common ancestor. The sequence *C* is determined by the parsimony criterion, i.e., the amino acid at position *p* of ancestor sequence *C* is the common amino acid at position *p* of the three sequences. For example, if amino acids at the first position of three sequences are ‘V’, ‘T’ and ‘T’, we assign ‘T’ as the amino acid at the first position of their common sequence *C*. If amino acids of three sequences are all different, unknown amino acid ′ − ′ is assigned as their common amino acid. The distances from *C* to sequences *S*_1_, *S*_2_, and *S*_3_ are smaller than that between the sequences. The pairwise amino acids in three sequence pairs (*S*_1_, *C*), (*S*_2_, *C*) and (*S*_3_, *C*) are counted into another 20 × 20 frequency matrix *F*_3_. We also randomly choose *N* triplets to create the summary statistics of *F*_3_. For each alignment, 400 relative rates of change of the *F*_3_ matrix summarizing amino acid substitutions from the alignment are used for training deep learning models.

##### Deep learning architecture

We used the ResNet-18 architecture (He et al. 2016) to train two networks: the pModelDetector network with 400 summary statistics from the Pairwise extraction method and the ModelDetector network with 800 summary statistics from both Pairwise and Triplet extraction methods (see Fig. 2). We reshaped 400 summary statistics and 800 summary statistics into shapes of 1 × 20 × 20 and 1 × 20 × 40, respectively. We used ReLu activation function for ResNet blocks and softmax activation function for the last layer. For gradient descent we used Adam optimizer (Kingma and Ba 2015). The label of a simulation alignment is the model used to generate the alignment. Our deep learning networks will assign an input alignment with one of the 9 labels: Q.plant, Q.bird, Q.yeast, Q.mammal, Q.insect, Q.pfam, LG, WAG or JTT.

**Fig. 2.**
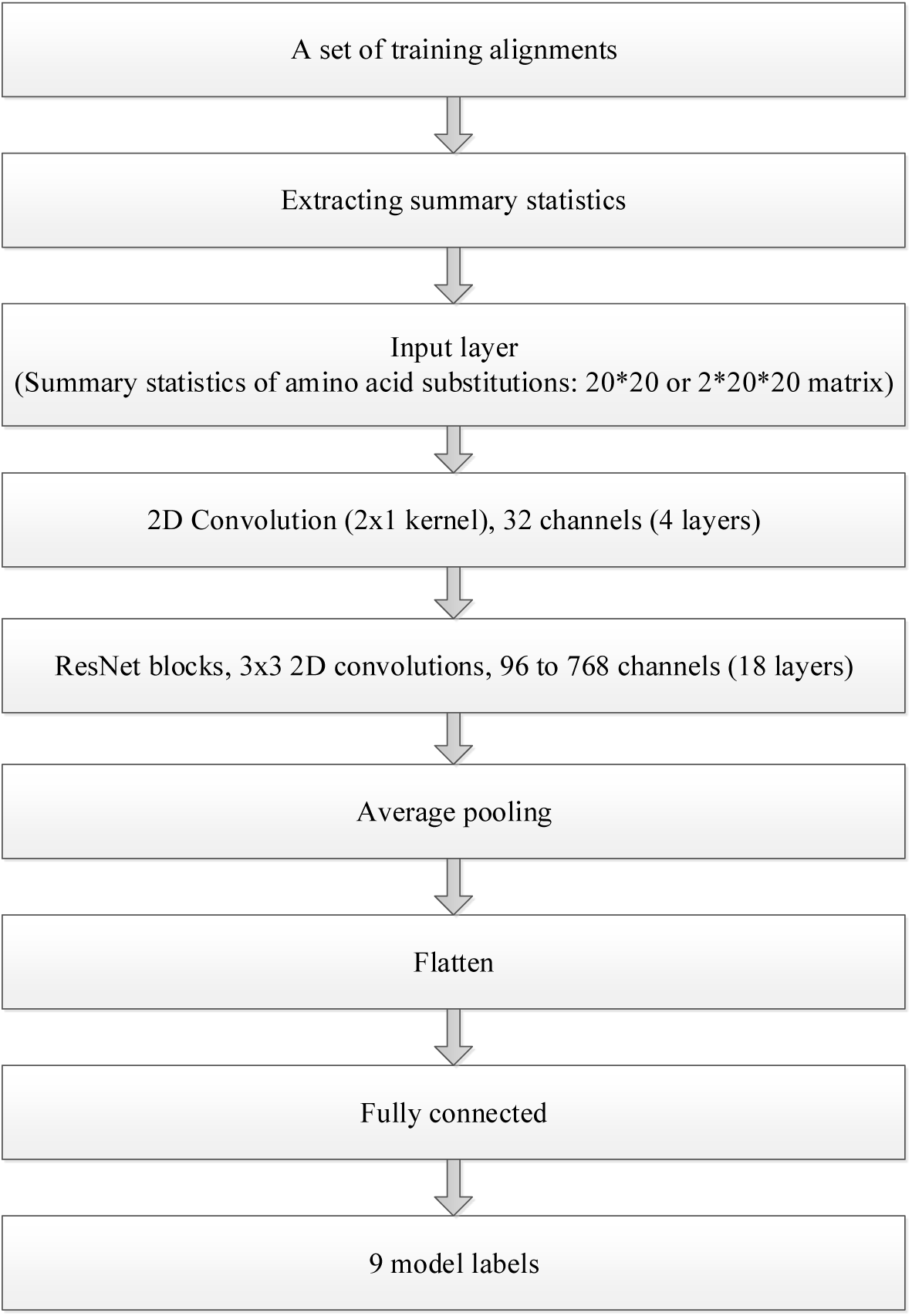
The deep learning architecture based on ResNet-18 to train two networks from a set of protein alignments.

We used 64,800 training alignments to train pModelDetector and ModelDetector networks using 30 epochs and a batch size of 40 (we tried more epochs but did not get better results). We used Tensorflow 2.13.1 (Ghemawat et al. 2016) to train the networks on a computer with 8 cores (no GPU required) and set 20% of training alignments for network validation.

For investigating the overfitting aspect, we performed our experiments five times. For each run, the real alignments were randomly separated into the training part and the testing part to simulate training and testing alignments. The networks were trained and tested from the simulation alignments. The average accuracy and the variance were calculated from the five runs to evaluate the overfitting problem.

## Results

### Summary statistics analysis

First, we analyzed the distribution of summary statistics calculated from simulation data. For each summary statistic, we calculated its variance from simulation alignments for different models. We used the boxplot to display variances and standard deviations of all summary statistics for the 9 models (see Fig. 3). Most of summary statistics have low variances (only few summary statistics have variances greater than 0.00015). The distributions of variances across different models exhibit similarity. We also observe that the variances of summary statistics calculated by the Pairwise and Triplet extraction methods are similar.

**Fig. 3.**
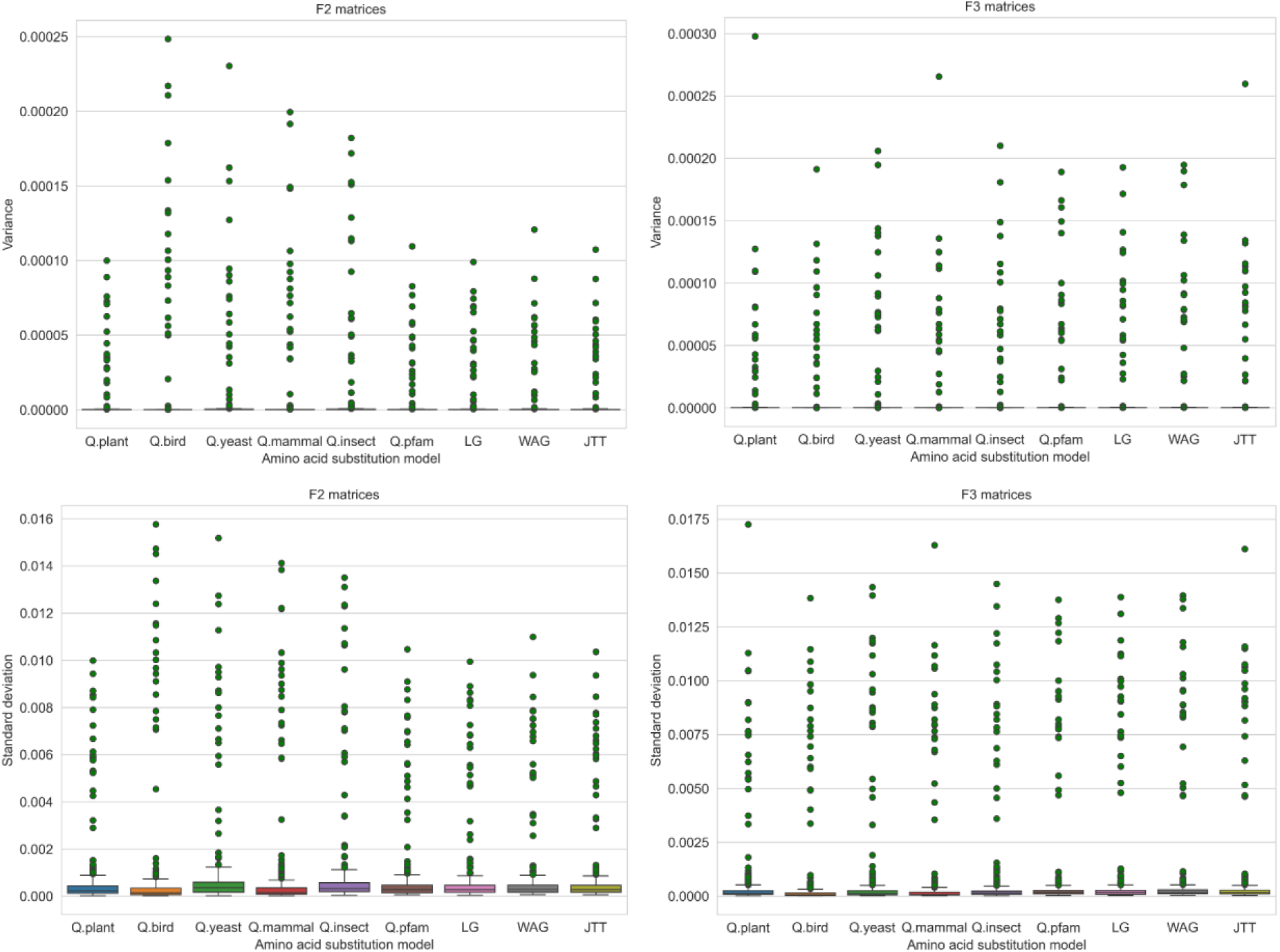
The variance and standard deviation of summary statistics F2 and F3 calculated from simulation alignments.

We also investigated the correlations between summary statistics extracted from simulated data and those from real data. For each clade-specific simulation alignment *D*, we calculated the Pearson correlation between F2 matrix (and F3 matrix) obtained from *D* and that from the real alignment used to simulate *D*. Fig. 4 shows that summary statistics obtained from the simulation data and those from the real data are highly correlated. The average correlation of F2 matrices ranges from 0.88 (yeast) to 0.94 (mammal). The correlations of more than 95% of alignments are higher than 0.8.

**Fig. 4.**
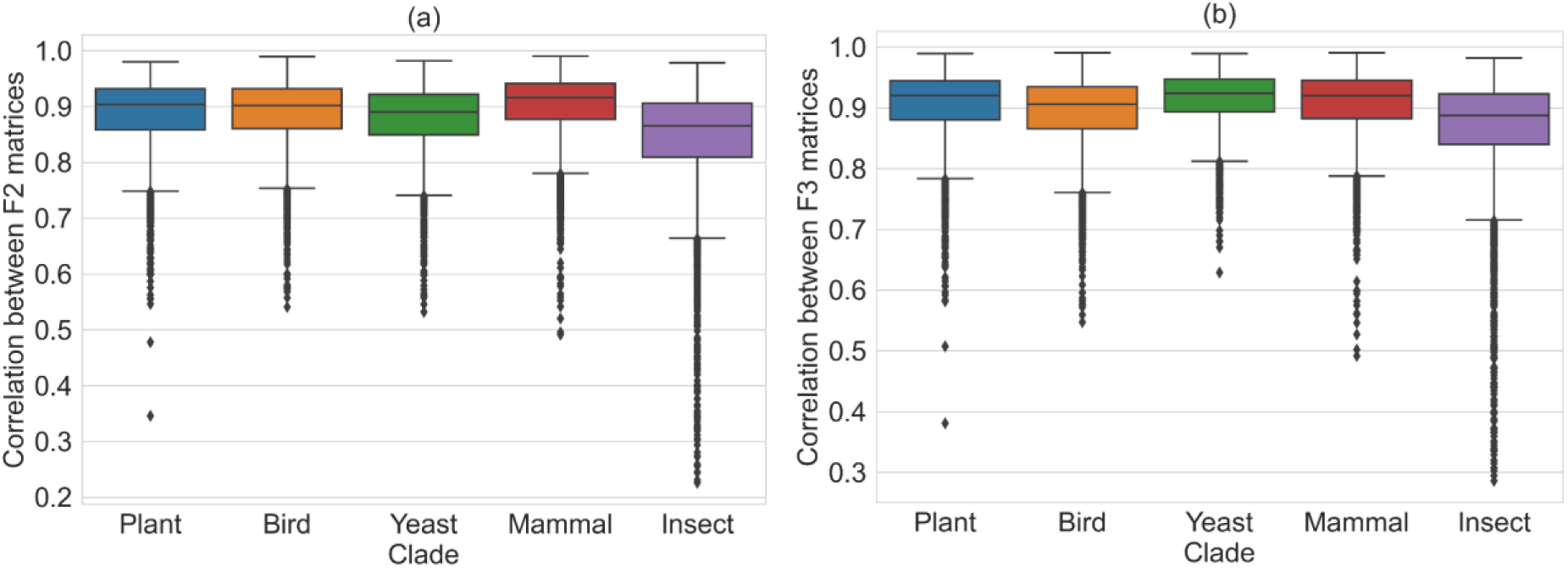
The correlation between F2 matrices (a) and F3 matrices (b) of real data and simulation data.

Finally, we evaluated the correlations between F2 and F3 matrices which were extracted from the same simulation alignment for all 9 models (see Fig. 5). The correlations between F2 and F3 matrices are greater than 0.9 for most alignments except some alignments with Q.yeast and Q.insect models. The average correlation between the F2 and F3 matrices is about 0.98.

**Fig. 5.**
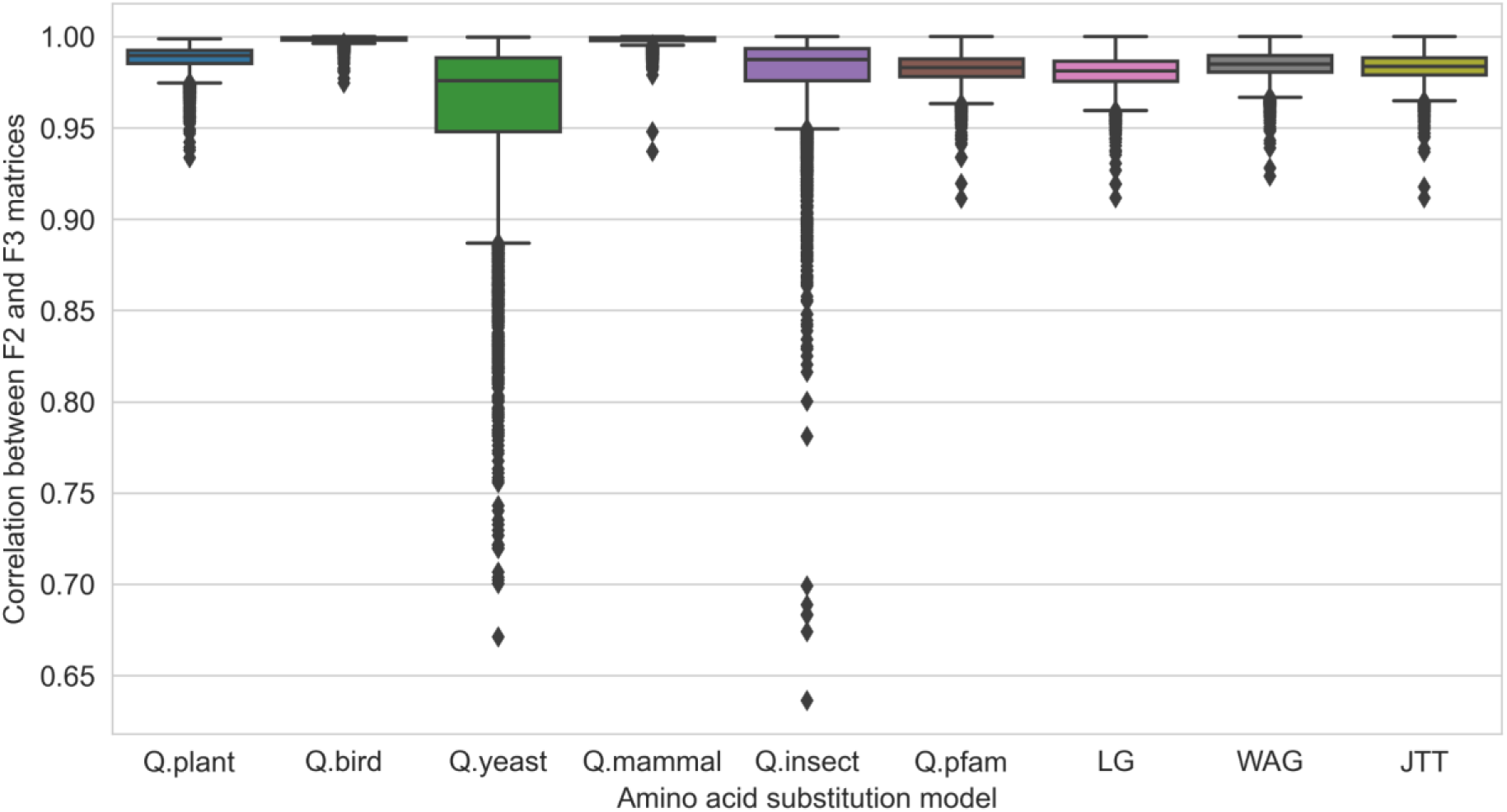
The correlations between the F2 and F3 matrices extracted from the same alignment.

### Network evaluation

First, we examined the accuracy improvement of our networks during the training process. Fig. 6 shows the training and validation accuracy (loss) of pModelDetector and ModelDetector networks. ModelDetector had slightly higher validation accuracy (the percentage of validation alignments correctly inferred) than pModelDetector. Both training and validation accuracy of the two networks increased to more than 0.95 after a few epochs.

**Fig. 6.**
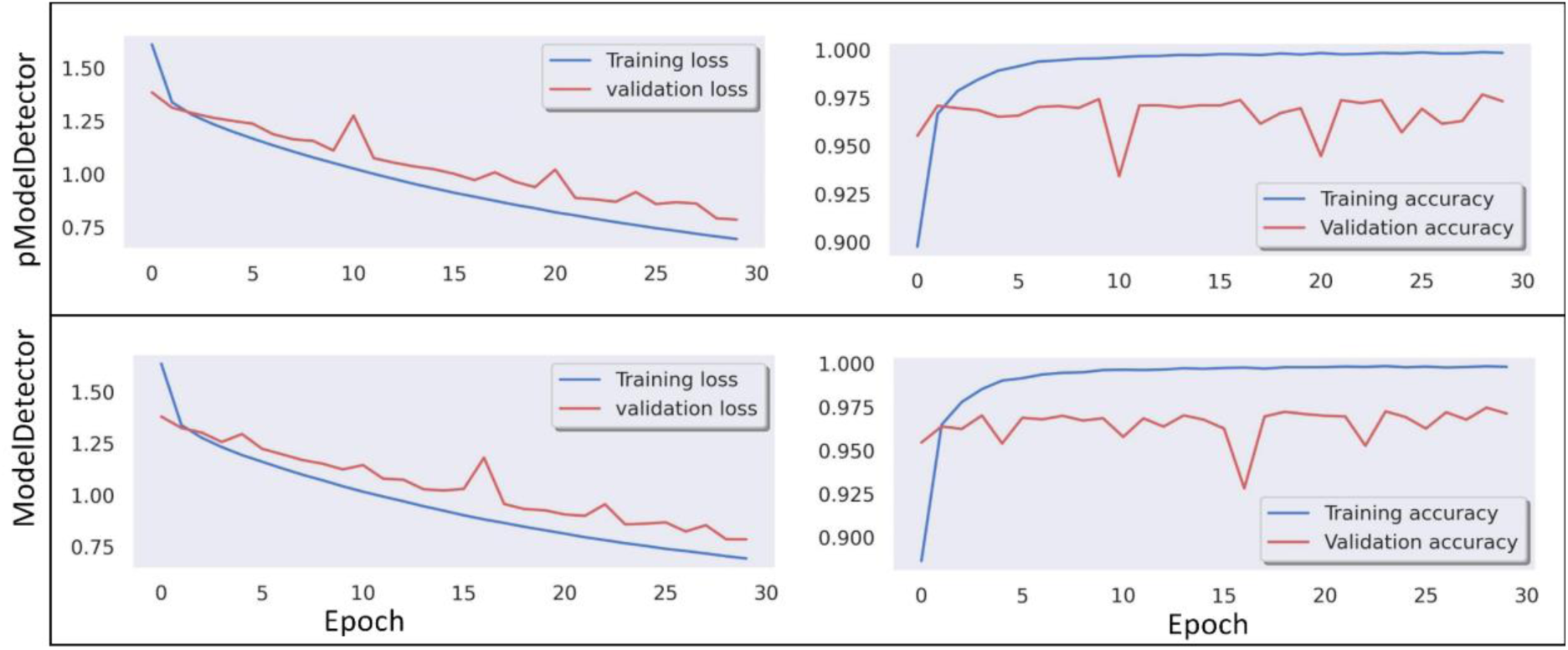
The training and validation accuracy of pModelDetector and ModelDetector networks during the training process.

Second, we analyzed the performance of pModelDetector and ModelDetector networks in comparison with that of the maximum likelihood method ModelFinder on testing alignments (see Fig. 7). ModelFinder was restricted to select the best model from a set of 9 models as used in the deep learning networks. The average accuracy (the percentage of testing alignments correctly inferred) of pModelDetector, ModelDetector, and ModelFinder was 96.78%, 97.45%, and 97.88%, respectively. ModelFinder was slightly better than the deep learning networks; while ModelDetector outperformed pModelDetector. The accuracy of all methods got higher with longer alignments. For example, the accuracy of all methods was only around 90% for alignments with 100 sites; but increased to more than 99% for 2000-site alignments (i.e., 99.1% for pModelDetector; 99.6% for ModelDetector; and 100% for ModelFinder).

**Fig. 7.**
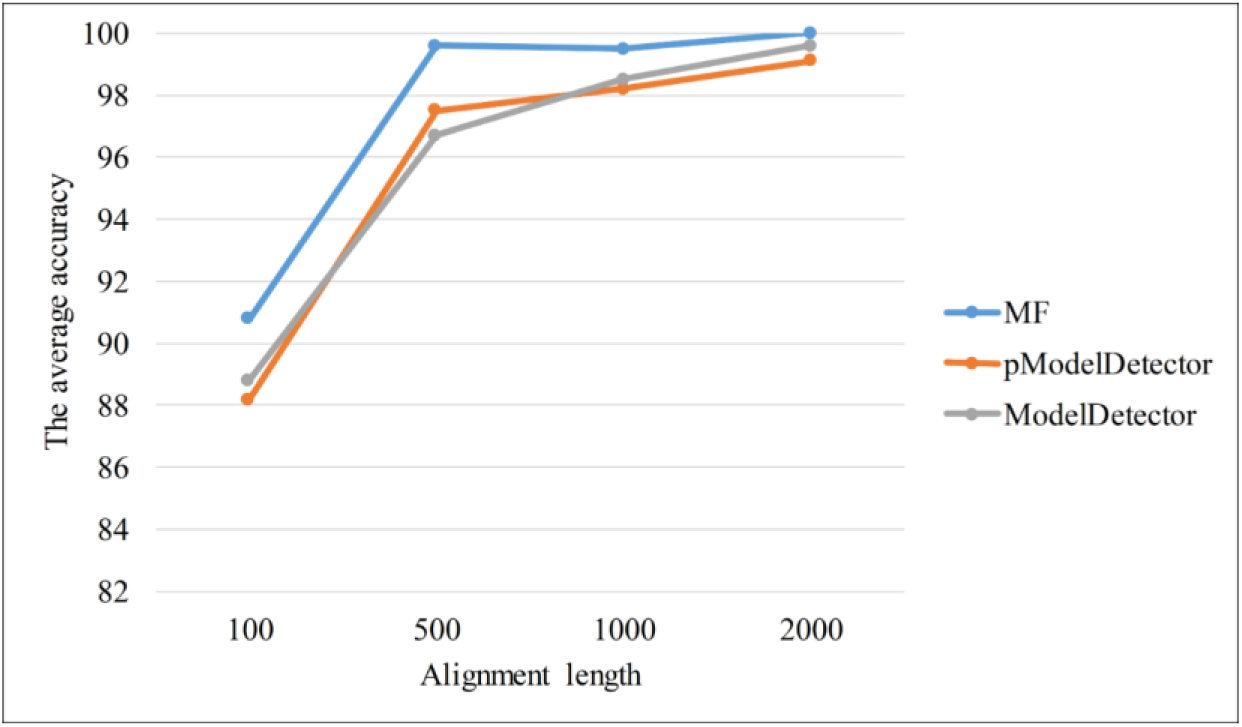
The average accuracy of ModelFinder (MF), pModelDetector and ModelDetector on testing alignments.

Fig. 8 shows the accuracy of all methods for different models. ModelFinder had high accuracy of > 96% for 8 out of 9 models except Q.bird (87.92%). A number of Q.bird alignments were wrongly predicted as Q.mammal alignments due to the high correlation between Q.bird and Q.mammal models (i.e., their Pearson correlation of 0.98). ModelDetector also had high accuracy of > 95% for 8 models except LG. Some small LG alignments were incorrectly assigned as Q.pfam alignments because both LG and Q.pfam models were estimated from the same dataset (the Pearson correlation of 0.99).

**Fig. 8.**
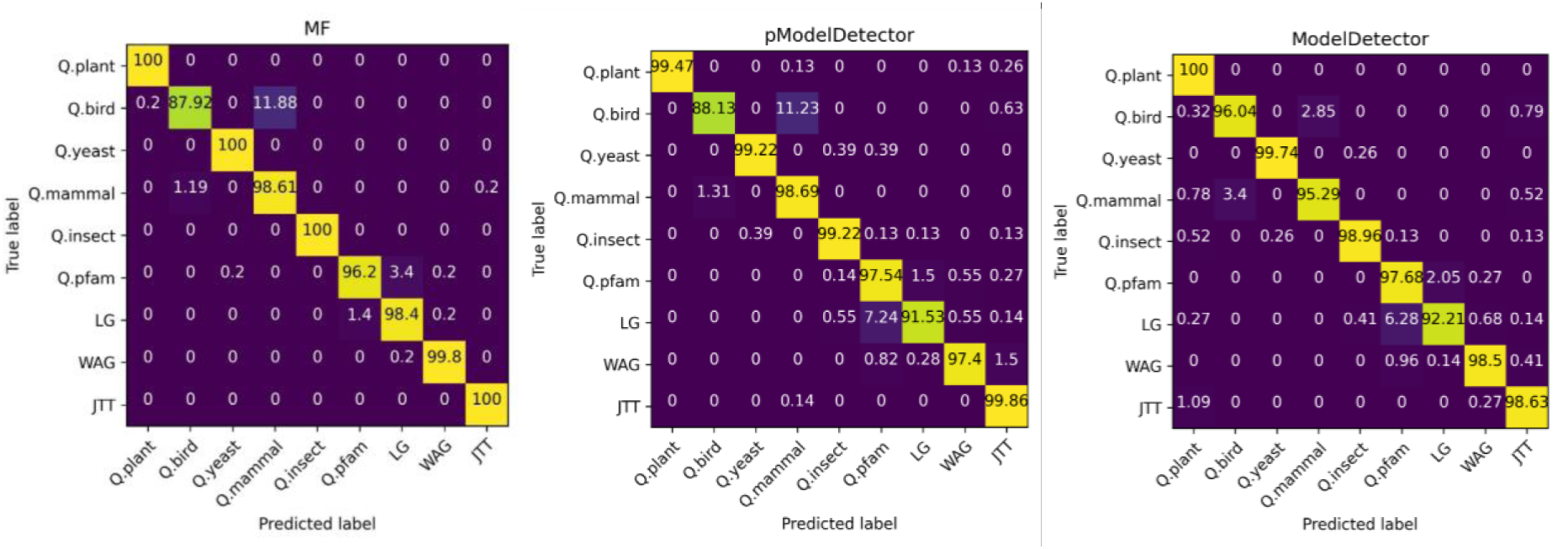
Confusion matrices of ModelFinder (MF), pModelDetector, and ModelDetector on testing alignments.

#### Overfitting examination

The pModelDetector and ModelDetector networks were trained and tested on five different datasets (see Table 2). The accuracy of both networks was high (> 96%) and stable (standard variation ≤ 0.5) for all datasets. The average accuracy of pModelDetector and ModelDetector over five datasets was 97% and 97.1%, respectively. The accuracy of ModelDetector for five datasets ranges from 96.2% to 97.5%. The performance of the networks across models were also stable (the smallest standard deviation of 0.2% for Q.plant; and the greatest standard deviation of 4.0% for Q.pfam).

**Table 2.** The accuracy of pModelDetector and ModelDetector on five different datasets.

|  |  | Dataset1 | Dataset2 | Dataset3 | Dataset4 | Dataset5 | Mean (standard deviation) |
| --- | --- | --- | --- | --- | --- | --- | --- |
| pModelDetector | Q.plant | 99.5 | 99.7 | 99.5 | 99.7 | 99.2 | 99.5 (0.2) |
|  | Q.bird | 88.1 | 93.8 | 98.2 | 93.1 | 91.5 | 92.9 (3.3) |
|  | Q.yeast | 99.2 | 98.8 | 99.0 | 99.6 | 99.7 | 99.2 (0.3) |
|  | Q.mammal | 98.7 | 96.3 | 93.2 | 94.2 | 97.9 | 96.1 (2.1) |
|  | Q.insect | 99.2 | 98.7 | 99.5 | 99.7 | 97.8 | 98.9 (0.7) |
|  | Q.pfam | 97.5 | 95.5 | 96.1 | 96.3 | 96.3 | 96.3 (0.7) |
|  | LG | 91.5 | 94.6 | 91.8 | 94.3 | 91.1 | 92.7 (1.5) |
|  | WAG | 97.4 | 98.4 | 99.1 | 99.5 | 98.0 | 98.5 (0.7) |
|  | JTT | 99.9 | 97.6 | 99.3 | 98.1 | 97.4 | 98.5 (1.0) |
|  | **Average** | **96.8** | **97.0** | **97.3** | **97.2** | **96.5** | **97 (0.3)** |
| ModelDetector | Q.plant | 100 | 99.9 | 99.7 | 99.4 | 99.5 | 99.7 (0.2) |
|  | Q.bird | 96 | 96.4 | 96.3 | 91.1 | 91.3 | 94.2 (2.5) |
|  | Q.yeast | 99.7 | 99.3 | 98.9 | 99.9 | 99.8 | 99.5 (0.4) |
|  | Q.mammal | 95.3 | 92.6 | 95.0 | 99.4 | 97.1 | 95.9 (2.2) |
|  | Q.insect | 99.0 | 98.5 | 98.9 | 94.9 | 99.5 | 98.2 (1.6) |
|  | Q.pfam | 97.7 | 87.3 | 96.9 | 97.7 | 93.7 | 94.7 (4.0) |
|  | LG | 92.2 | 98.1 | 94.5 | 98.3 | 93.3 | 95.3 (2.5) |
|  | WAG | 98.5 | 95.4 | 98.3 | 98.9 | 98.7 | 97.9 (1.3) |
|  | JTT | 98.6 | 98.4 | 98.5 | 96.3 | 97.6 | 97.9 (0.9) |
|  | **Average** | **97.5** | **96.2** | **97.4** | **97.3** | **96.7** | **97.1 (0.5)** |

#### Time analyses

The pModelDetector and ModelDetector networks were trained on a computer with 8 cores and no GPU required. The pModelDetector network required 801 seconds per epoch. ModelDetector has a double number of summary statistics, therefore it required more training time than pModelDetector (i.e., 1412 seconds per epoch). Totally, it took about 7 hours to train the pModelDetector network; and about 12 hours to train the ModelDetector network from 64,800 alignments. The training time of ModelDetector network increased proportionally with the number of training alignments.

We evaluated the inferring time (the total time for summary statistics creation and model prediction) of the methods on testing alignments (see Fig. 9). The ModelDetector and pModelDetector networks were much faster than ModelFinder for all cases. The inferring time of ModelFinder raised rapidly with an increase in alignment length or the number of taxa. For simulation alignments, the average running time of ModelFinder was 2.2, 4.6, 8.1 and 14.8 seconds for alignments with 100, 500, 1000 and 2000 sites, respectively. The inferring time of both pModelDetector and ModelDetector networks increased slightly with the increase of alignment length. For example, it required 1.39 seconds for an alignment with 100 sites; and 1.40 seconds for a 2000-site alignment.

**Fig. 9.**
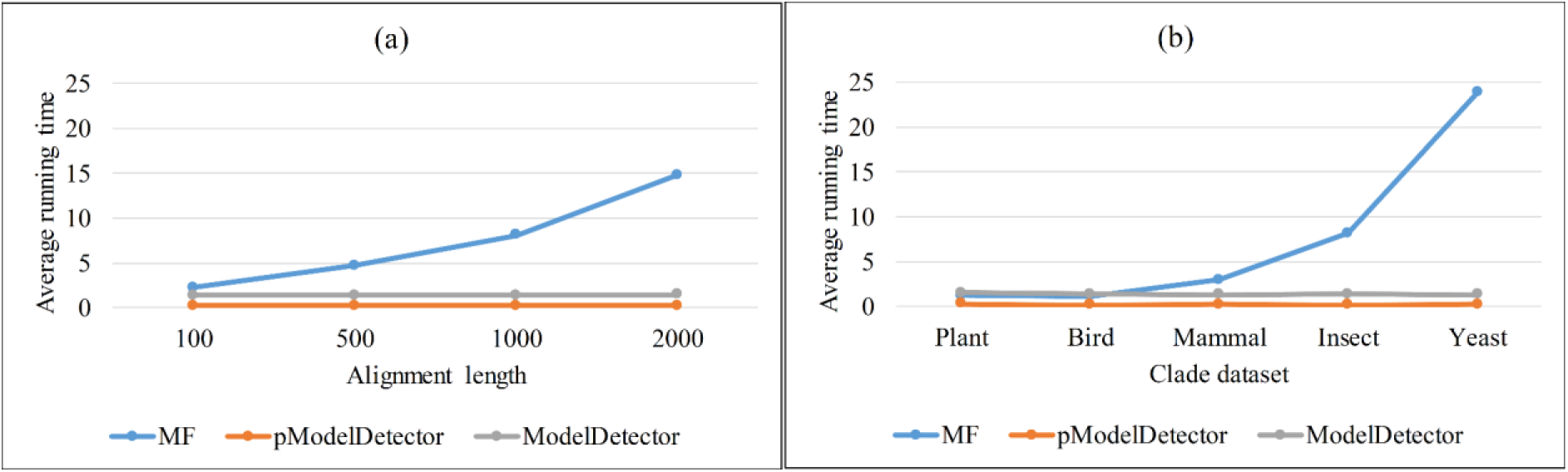
The average inferring time (in seconds) of ModelFinder (MF), pModelDetector and ModelDetector networks per alignment.

For clade-specific alignments, ModelFinder took an average of 1.2 seconds to select the best-fit model for a 35-taxa plant alignment; and increased to 23.8 seconds for a 332-taxa yeast alignment. The inferring time of the deep learning networks were very low for all alignments. For example, the average inferring time of ModelDetector was 1.31 seconds for a 35-taxa plant alignment; and 1.34 seconds for a 332-taxa yeast alignment.

Finally, we examined the running time of ModelDetector and ModelFinder on large simulation alignments with 50 taxa and alignment lengths from 1000 sites to 1,000,000 sites (∼3000 plant genes or 1500 bird genes). Table 3 shows that ModelDetector required 82.8 seconds for an alignment with 500,000 sites or 186.1 seconds for an alignment with 1,000,000 sites. ModelFinder required ∼5,733 seconds to select the best-fit model for an alignment with 500,000 sites or ∼12975 seconds for an alignment with 1,000,000 sites.

**Table 3.** The inferring times (in seconds) of ModelDetector and ModelFinder on alignments with 50 taxa and lengths from 1,000 to 1,000,000 sites.

| Length | ModelDetector |  |  | ModelFinder |
| --- | --- | --- | --- | --- |
|  | Summary statistics creation | Model Prediction | Total |  |
| 1,000 | 1.3 | 1.8 | 3.1 | 2.1 |
| 10,000 | 1.6 | 1.9 | 3.5 | 26.2 |
| 50,000 | 8.6 | 1.9 | 10.5 | 182.2 |
| 100,000 | 16.3 | 1.9 | 18.2 | 432.5 |
| 500,000 | 82.8 | 1.9 | 84.7 | 5733.3 |
| 1,000,000 | 184.2 | 1.8 | 186 | 12975.4 |

## Discussion

The maximum likelihood method has been widely used to select the best fit models for alignments based on the information criteria such as BIC score. However, theoretical concerns about the suitability of information criteria in selecting proper models have been raised in different studies. Moreover, the computational burden prevents the maximum likelihood method to work with long alignments or alignments with many species.

Recently, the deep learning network ModelRevelator has been proposed for nucleotide model selection. In this paper, we introduced another deep learning network ModelDetector for amino acid model selection. Experiments on simulation data showed that the accuracy of ModelDetector was comparable with that of the likelihood-based method ModelFinder. Both ModelRevelator and ModelDetector networks are based on the convolutional neural network ResNet-18 architecture. The key difference between ModelRevelator and ModelDetector is the strategy to extract summary statistics from alignments. ModelRevelator extracts 260,000 summary statistics from a nucleotide alignment while ModelDetector calculates only 800 summary statistics from a protein alignment. The ModelRevelator network was trained with 242,688 alignments that required powerful computers with GPU; while the ModelDetector network can be trained with 68,000 alignments on a personal computer without GPU. For inference, both ModelRevelator and ModelDetector are very fast and applicable on personal computers. They are much faster than the maximum-likelihood method, especially for large alignments. ModelDetector can detect the amino acid model for a 1,000,000-site protein alignment in a few minutes.

As we do not know ‘true’ models of real alignments, all deep learning networks were trained and evaluated on simulation data. It is always a challenge for alignment simulators to create alignments that can resemble real alignments. We used the AliSim program to simulate alignments from real alignments, i.e., trees and parameters of site rate models were estimated from real alignments. The mixture models were not included in this study because they require more complicated extraction methods. As real alignments might be more complex than simulation alignments (e.g., different parts of a real alignment might follow different known or unknown models), the lack of real alignments with ‘true’ models is a challenge for the deep learning networks.

We also explored the extension of the ModelDectector network to more models. To this end, we trained ModelDetector13 network with 13 models (including four additional general models VT, PMB, Blosum62, and Dayhoff) from 93,600 training alignments. Experiments showed that the accuracy of ModelDetector13 with 13 models was about 97.9% (see Appendix) that was similar to that of ModelDetector with 9 models. The training time of our networks increased proportionally with the number of training alignments (i.e., 12 hours for training ModelDetector with 68,400 alignments; and 15 hours for training ModelDetector13 with 93,600 alignments). The inference times of ModelDetector and ModelDetector13 networks were similar and not affected by the number of models.

The accuracy of ModelDetector got higher when trained with more alignments, i.e., 78.1%, 91.1%, and 97.5% for the cases of 18,000; 32,400; and 64,800 training alignments, respectively. ModelDetector was trained and tested on alignments with at least 100 sites because it requires a sufficient number of amino acid substitutions to infer correct models. The accuracy of ModelDetector got higher with longer alignments. It assigned correct models for more than 99% of 2000-site alignments. The accuracy of ModelDetector dropped below 90% for 100-site alignments, indicating that it is not advisable for use with shorter alignments.

## Author Contributions

N.H.T carried the experiments and drafted the manuscript. L.S.V conceptualized the study and wrote the final manuscript.

## Data Availability

The models, datasets and scripts used in this paper are available at https://doi.org/10.6084/m9.figshare.24711765.v14

## Statements and Declarations

### Appendix

**Fig. S10.**
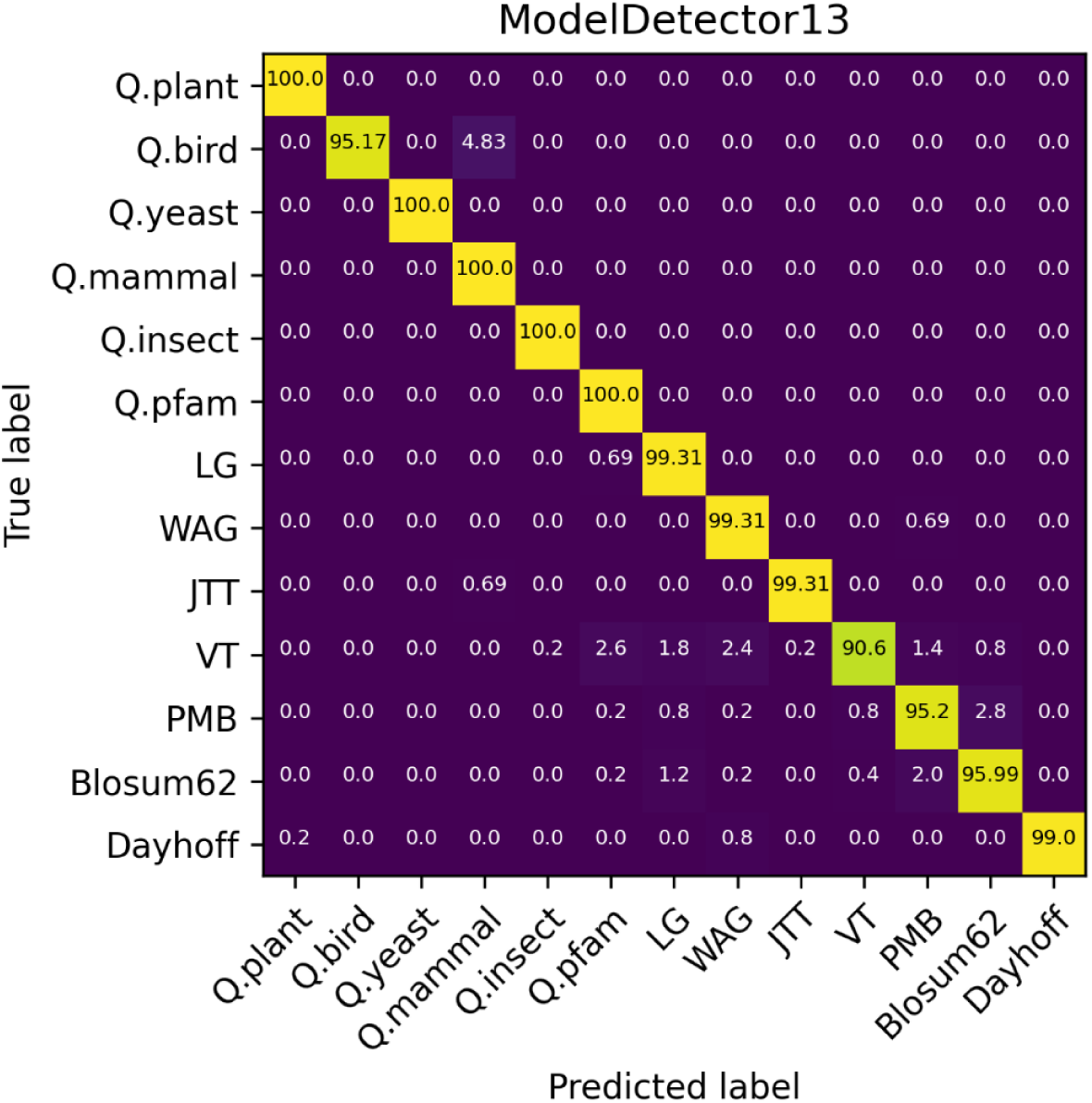
The confusion matrix of ModelDetector network trained with 13 models from 93,600 training alignments

